# Fast and cloning-free CRISPR/Cas9-mediated genomic editing in mammalian cells

**DOI:** 10.1101/612192

**Authors:** Paul T. Manna, Luther J. Davis, Margaret S. Robinson

**Affiliations:** Cambridge Institute for Medical Research, University of Cambridge, Cambridge Biomedical Campus, Cambridge, CB2 0XY, UK

**Author notes:** Corresponding author: Dr. Paul T. Manna.

## Abstract

CHoP-In (CRISPR/Cas9-mediated, Homology-independent, PCR-product Integration) is a fast and cloning-free strategy for genomic editing of mammalian cells. The desired integration fragment is produced as a PCR product, flanked by the Cas9 recognition sequences of the target locus. When co-transfected with the cognate Cas9/guide RNA, double strand breaks are introduced at the target genomic locus and at both ends of the PCR product. This allows incorporation into the genomic locus via hon-homologous end joining. The approach is versatile, allowing N-terminal, C-terminal or internal tag integration and gives predictable genomic integrations, as demonstrated for a selection of key membrane trafficking proteins. The lack of any donor vectors offers advantages over existing methods in terms of both speed and hands-on time. As such this approach will be a useful addition to the genome editing toolkit of those working in mammalian cell systems.

## Introduction

Whilst offering clear advantages over ectopic expression, endogenous protein tagging carries an inherent cost in both time and resources which can prove limiting in its application. With this in mind, we set out to develop a cloning-free approach, reducing the time and effort required to generate endogenous knock-ins via the CRISPR/Cas9 system.

The CRISPR/Cas9 system has radically simplified genome editing in mammalian cells (Jinek et al., 2012; Cho et al., 2013; Mali et al., 2013). Knockouts can now be created rapidly and easily, by relying on error-prone non-homologous end joining (NHEJ) to generate insertions or deletions (indels). Furthermore, short sequences such as small linear epitope tags can be introduced quickly and easily by homology-directed repair (HDR), using readily synthesised single-stranded oligodeoxynucleotide (ssODN) donors with short homology arms of 50-70 bp flanking the double strand break (DSB) site (González et el., 2014). Although fast, this approach does not allow incorporation of longer sequences such as fluorescent protein tags and thus requires the screening of a number of single cell derived colonies. The creation of larger knock-ins via homology-directed repair (HDR) remains relatively laborious, requiring time-consuming cloning of a donor vector with the integration cassette flanked by 0.5-1.5 kb homology arms. This is particularly problematic due to the widespread use of fluorescent protein tags to determine localisation and perform affinity isolation. There is clearly a need to simplify the process of fluorescent protein tag integration, particularly when aiming to tag multiple loci for example when attempting to validate the results of a screen.

NHEJ, the predominant repair mechanism in mammalian cells, is increasingly being investigated as an alternative route to producing large genomic knock-ins (Maresca et al., 2013, Lackner et al., 2015, He *et al.*, 2016, Schmid-Burgk et al., 2016, Suzuki et al., 2016, Sawatsubashi et al., 2017). For NHEJ mediated editing, the desired integration fragment is generally provided in a plasmid vector, flanked by Cas9 recognition sites for liberation once inside the target cell. The released fragment is then integrated, during the repair of a Cas9-mediated genomic DSB, via the NHEJ pathway. The reliance upon a donor vector imposes some limitations on these approaches. Generic NHEJ approaches use a small number of donor vectors for all knock-ins which greatly reduces the cloning required but also severely limits control of the final nucleotide sequence around the edit site (Lackner et al., 2015, Suzuki et al., 2016, Sawatsubashi et al., 2017). Conversely, more bespoke NHEJ approaches, those allowing greater control over the integrated DNA fragment, require individual donor vectors to be prepared for each knock-in (Schmid-Burgk et al., 2016). Therefore, while potentially much faster, NHEJ approaches are currently limited in terms of versatility. Removing the reliance upon a donor vector has the potential to combine the speed of generic NHEJ approaches with the versatility of bespoke approaches. However, previous attempts to employ a PCR product or restriction fragment as the donor have been unsuccessful, suggested to be due to a requirement for intracellular cleavage and co-processing to target the donor fragment for successful NHEJ integration (Lackner et al., 2015, Sawatsubashi et al., 2018).

Here we demonstrate a donor vector independent, NHEJ-mediated genome editing strategy for mammalian cells. Our methodology, which we term CHoP-In (**C**RISPR-mediated, **Ho**mology-independent, **P**CR-product **In**tegration), employs a PCR-generated donor, flanked by gene-specific guide RNA (gRNA) and protospacer adjacent motif (PAM) sequences incorporated via the PCR primers. Thus, donors can be produced quickly and easily, without the need for cloning. To demonstrate the utility, versatility and fidelity of our approach we have generated and characterised multiple HeLa cell lines expressing fluorescent protein fusions from their endogenous loci. Four example fusion-proteins are presented, each requiring a different site of tag integration and each showing a distinct but well characterised localisation within the endomembrane system.

## Results

### Requirements for CHoP-In genome editing

Two components need to be prepared for each CHoP-In experiment. The first is a plasmid (e.g. pX330) encoding Cas9 and a gRNA targeting the gene of interest. The second is a PCR product containing the desired integration fragment (e.g. a fluorescent protein tag), flanked by Cas9 recognition sites corresponding to the target gRNA and PAM sequences of the gene of interest, but in the reverse orientation. Thus, for a gRNA targeting the sense strand, the integration fragment must be flanked by gRNA and PAM sequences in the antisense orientation (Figure 1), and vice versa. This is to avoid reconstituting the genomic Cas9 target site. It also has the advantage of conferring directionality, as undesired reverse integrations will be excised via further CRISPR/Cas9 cleavage. Co-transfection of the plasmid and PCR product leads to cleavage of both the genomic DNA and the donor, which is then integrated into the genomic DSB by NHEJ (Figure 1).

**Figure 1.**
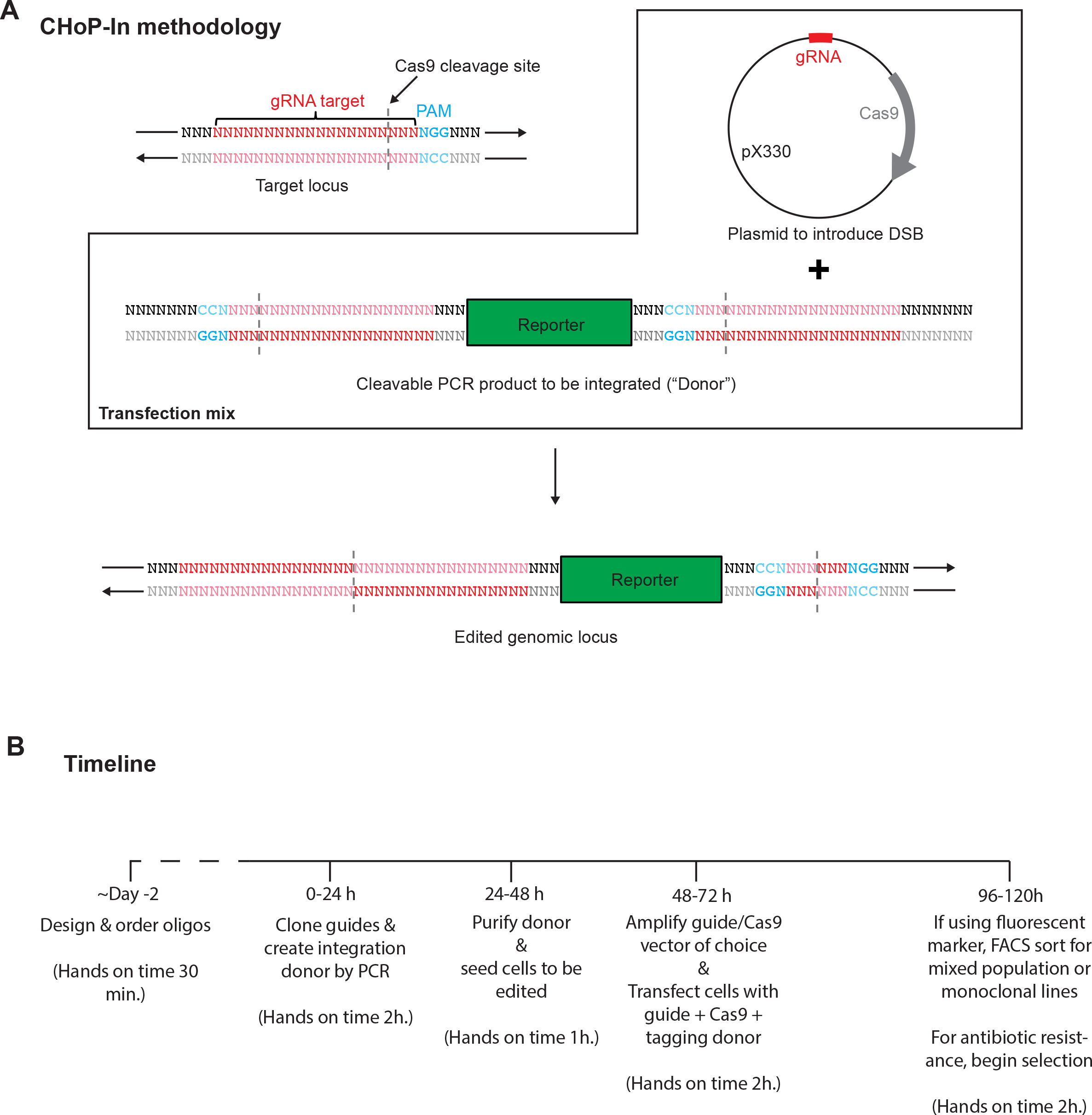
Genome editing by CHoP-In. CHoP-In genome editing relies utilizes NHEJ-mediated integration of a PCR-generated cassette into a CRISPR/Cas9-induced DSB. (A) To achieve this, two constructs must be prepared after identifying the genomic gRNA and PAM site. To introduce a DSB at the desired locus, a vector such as pX330 encoding both the gRNA and the Cas9 nuclease is constructed. Additionally, a CHoP-In integration cassette is produced by PCR, consisting of the desired integration fragment flanked by the same gene specific gRNA and PAM sites in the PCR primers. Importantly, the gRNA and PAM sites flanking the integration cassette must be in the reverse orientation with respect to genomic locus as this prevents reconstitution and re-cleaving of the sites following integration. (B) The whole process can be completed in approximately one week, giving a mixed population of edited cells with minimal hands-on time when compared with HDR mediated approaches.

PCR-generation of donors allows for integration of different exogenous DNA sequences, addition of linker sequences and conservation of frame by simply modifying the PCR primers, without any requirement for creating new donor vectors. Removing the need for cloning greatly speeds up the process of genomic editing, both in absolute terms and particularly in hands-on time requirement, a significant benefit over existing approaches (Figure 1B).

### N-Terminal tagging of RAB5C

N-terminal tagging is the optimal configuration for CHoP-In in non-haploid cell types as selection of a gRNA recognition site upstream of the translation start codon avoids indel generation in the coding sequence of other alleles. In addition, for some proteins, such as Rab GTPases, the N terminus is the only position where a tag can be inserted without compromising function. As with all NHEJ knock-in approaches, CHoP-In will generate a small scar from the gRNA and PAM sequences, which forms part of the linker sequence between the target protein and the tag. When generating an N-terminal tag, this scar can be minimised by selecting a sense strand-targeting gRNA, ideally as close to the start codon as possible.

As a demonstration of N-terminal tagging using CHoP-In, an emerald fluorescent protein (EmGFP) tag was introduced to the N terminus of the early endosomal Rab GTPase Rab5C (Figure 2A). To generate a RAB5C integration donor, EmGFP DNA was PCR-amplified using CHoP-In primers (Figure 2B). Because a sense strand-targeting gRNA was selected, the forward primer contained the gRNA and PAM sequence in the antisense direction, followed by a Kozak sequence prior to the start codon of the EmGFP. The reverse primer also contained the gRNA and PAM sequence, in the antisense direction when expressed, as well as a flexible peptide linker (GGSGG) between the tag and RAB5C. The predicted edited protein sequence is shown in Figure 2B. HeLa cells were transfected with pX330:RAB5C (encoding the gRNA and Cas9), together with donor PCR product, either encoding a frame corrected EmGFP lacking gRNA recognition sequences or a full CHoP-In integration fragment as described in Figure 2B. 48 hours after transfection these cell populations were analysed and sorted by flow cytometry. Transfection of a PCR product encoding EmGFP but lacking gRNA recognition sequences led to little to no stable integration the EmGFP tag at the RAB5C locus. The percentage of positive cells following CHoP-In editing suggests a knock-in efficiency of approximately 0.5% (Figure 2C), in line with other NHEJ-mediated knock-in approaches (Lackner et al., 2015, He et al., 2016, Schmid-Burgke et al., 2016). Importantly, subsequent imaging of the mixed population of EmGFP-positive cells showed homogeneous subcellular distribution. Consistent with the known distribution of Rab5 proteins (Munro, 2004), much of the EmGFP localised to early endosomes, marked by Alexafluor-555 labelled transferrin that had been endocytosed for 15 minutes (Figure 2D). Western blotting of the mixed population of edited cells with an antibody against RAB5C showed a band at the expected weight for a correctly tagged allele (Figure 2E).

**Figure 2.**
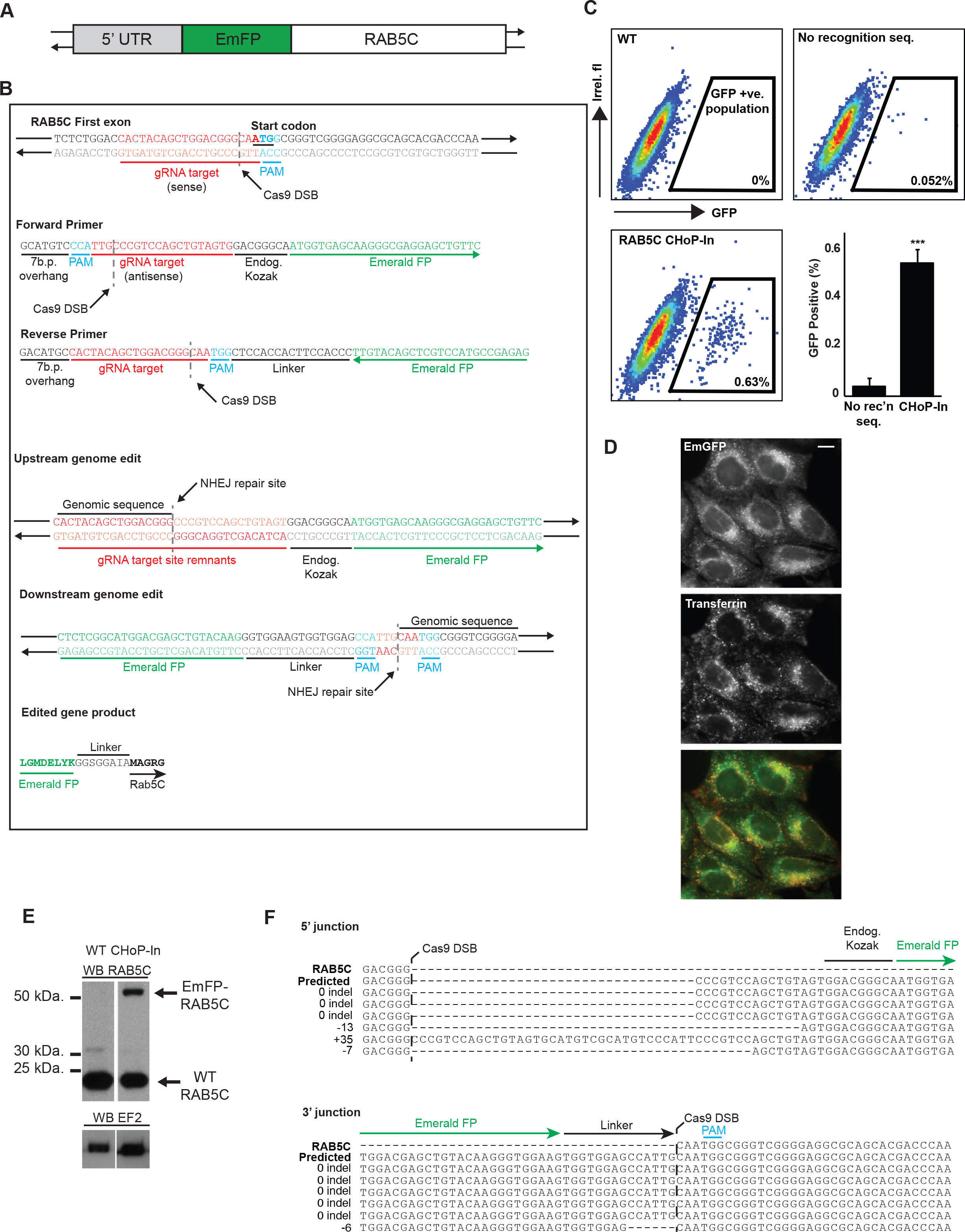
Generation and characterisation of an EmGFP-RAB5C HeLa line. (A) Sequence encoding an N-terminal EmGFP tag was integrated into the endogenous RAB5C locus by CHoP-In. (B) A sense strand gRNA and PAM site was selected immediately upstream of the RAB5C start codon and CHoP-In primers were designed to amplify DNA encoding EmGFP, flanked by the necessary gRNA and PAM sites for intracellular cleavage and NHEJ-mediated integration. (C) Following transfection with px330-RAB5C together with PCR donor fragments consisting of a frame corrected EmGFP tag without any Cas9 recognition site (no recognition seq.) or a full CHoP-In donor fragment (ChoP-In), cells were analysed and sorted by flow cytometry. WT cells are untransfected HeLa. Data are shown as FACS plots from individual experiments as well as mean data (+/− s.d.) from three independent experiments. (D) Fluorescence microscopy revealed EmGFP signal (green) in cells from this mixed population which colocalised well with Alexafluor-555 labelled endocytic tracer transferrin (red), following uptake of the marker for 15 minutes in order to label early endosomes (scale bar equals 5 μm). (E) Immunoblotting with an antibody against RAB5C revealed the presence of a higher molecular weight band corresponding to the EmGFP-RAB5C fusion in lysates from the mixed population. (F) Sanger sequencing of integration junctions showed the fidelity of NHEJ mediated knock-in of EmGFP into the RAB5C locus.

To assess the fidelity of CHoP-In editing, Sanger sequencing was carried out on targeted loci PCR amplified from genomic DNA prepared from the mixed population. 8 of 12 sequenced junctions had no indel, and only one indel was seen at a 3’ junction, generating an in-frame deletion within the linker sequence between the EmGFP tag and Rab5C (Figure 2F). The observed fidelity of the CHoP-In approach thus compares well with that of other NHEJ-mediated knock-in approaches (Lackner et al., 2015, He et al., 2016).

### C-terminal tagging of ATP6V1G1

It is not always possible to tag a protein at its N terminus, for example where this tag placement would disrupt intermolecular interactions or interfere with sorting signals such as mitochondrial or endoplasmic reticulum signal sequences. In these cases, a C-terminal tag is the most common solution. The viability of C-terminal tagging by CHoP-In was demonstrated via the generation of HeLa cells expressing a C-terminally tagged ATP6V1G1, a subunit of the vacuolar ATPase localizing to late endosomes and lysosomes (Forgac, 2007) (Figure 3A). An antisense-oriented gRNA site just upstream from the stop codon was selected in order to minimise the integration scar (Figure 3B). EmGFP again served as the tag, flanked in the integration donor by gRNA and PAM sequences in the sense orientation (Figure 3B). Two days after transfection, EmGFP-positive cells were isolated by flow cytometry, this time accounting for approximately 2% of the population. Consistent with the known localisation of the ATP6V1G1 protein the EmGFP fluorescence in this mixed population co-localised well with the endolysosomal marker Magic Red, a cathepsin B substrate liberating a fluorescent cresyl violet dye upon hydrolysis and therefore marking catalytically active degradative compartments (Figure 3C) (Creasy et al., 2007, Bright et al., 2016). Immunoblotting with an antibody to ATP6V1G1 revealed a band at the expected molecular weight for the tagged protein in the mixed population, at approximately one third the intensity of wild-type ATP6V1G1 in unedited HeLa cells. (Figure 3D). This suggests that on average, one of the three ATP6V1G1 alleles in our mixed population was tagged.

**Figure 3.**
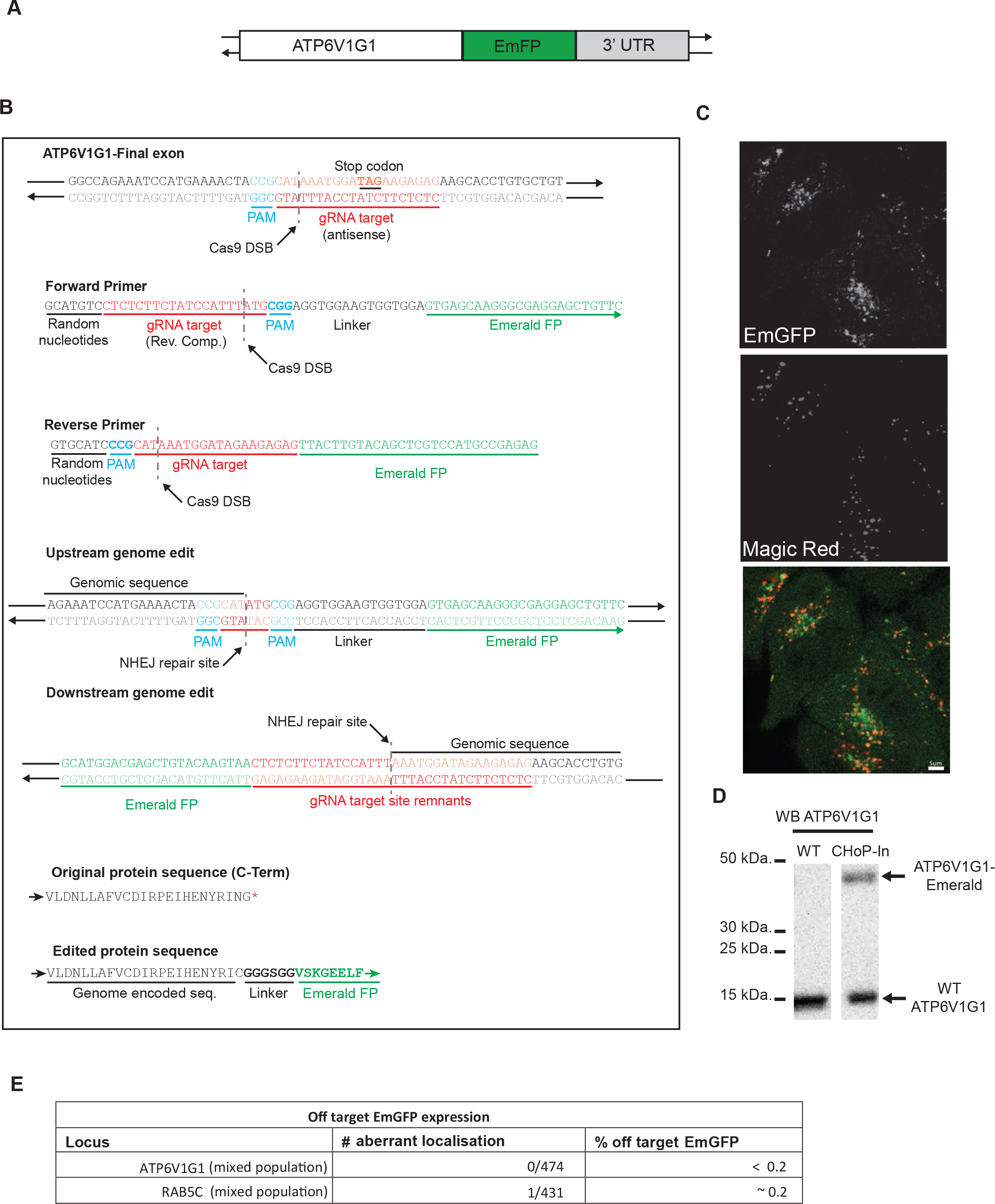
C-terminal tagging of ATP6V1G1. (A) ATP6V1G1 was tagged at its C terminus with EmGFP. (B) An antisense strand targeting gRNA and PAM site was selected which would create a DSB slightly upstream of the endogenous stop codon. EmGFP was amplified with CHoP-In primers encoding sense orientation gRNA and PAM sites. (C) Good colocalisation of EmGFP signal with the endolysosomal marker Magic Red was seen In a flow cytometry isolated EmGFP positive mixed population (scale bar equals 5 μm). (D) Immunoblotting with an antibody against ATP6V1G1 confirmed expression of higher molecular weight, EmGFP-tagged ATP6V1G1 from its endogenous locus. (E) Off target expression of EmGFP was assessed by examining mixed populations of cells for aberrant GFP localisation.

We next wanted to assess the reliability of localisation information obtained from CHoP-In edited cells. To do this we examined mixed populations of tagged cells for any examples of aberrant EmGFP localisation by microscopy. We found extremely low levels of aberrant EmGFP localisation suggesting that off target expression of tagging cassettes is not a major problem following CHoP-In genome editing (Figure 3E)

As opposed to N-terminal tagging, C-terminal tagging via CHoP-In has the potential to disrupt other alleles through NHEJ-mediated indel generation without tag integration. To explore this possibility monoclonal cell lines were generated from the ATP6V1G1-EmGFP mixed population. In one of these lines, clone 7, as well as an apparently correct EmGFP fusion protein, assessed by western blot and fluorescence imaging (Supplemental figure S1), an additional shifted band and was seen by western blot which was too small to represent EmGFP integration. Sequencing confirmed correct integration of the EmGFP into at least one allele, but also revealed an additional allele harbouring a one base pair indel creating a frameshift mutation extending the reading frame and accounting for the apparent molecular weight shift (Supplemental figure S1).

To assess the wider applicability of CHoP-In editing in other commonly used cell types we attempted to tag the RAB5C locus in HEK293-T cells and the ATP6V1G1 locus in NRK cells. In both cases we observed comparable results to those obtained in HeLa cells (Supplemental figure S2) suggesting that our approach is likely to work in a range of mammalian model cell systems.

### Internal tagging

As a further test of CHoP-In versatility, internal tags were introduced into two proteins for which both N and C-terminal tagging have been shown to be disruptive: subunits of the AP-1 and AP-2 adaptor protein complexes. These represent challenging targets as tag placement is constrained to regions of high intrinsic disorder between or within folded domains in both proteins. The detailed CHoP-In targeting strategy for these two genes is shown in supplemental figures S2-3. Mixed populations of HeLa cells edited at either exon 7 of the AP2M1 gene to express an internal EmGFP fusion (Figure 4A) (Hong et al., 2015), or exon 20 of the AP1G1 gene to express an internal mCherry fusion (Figure 4C) (Robinson et al., 2010) were generated. The subcellular distribution of fluorescence was homogeneous amongst the mixed cell populations and correctly localised, with AP2M1 showing discrete puncta at the plasma membrane (Figure 4B) and AP1G1 showing a tubulovesicular perinuclear distribution (Figure 4D).

**Figure 4.**
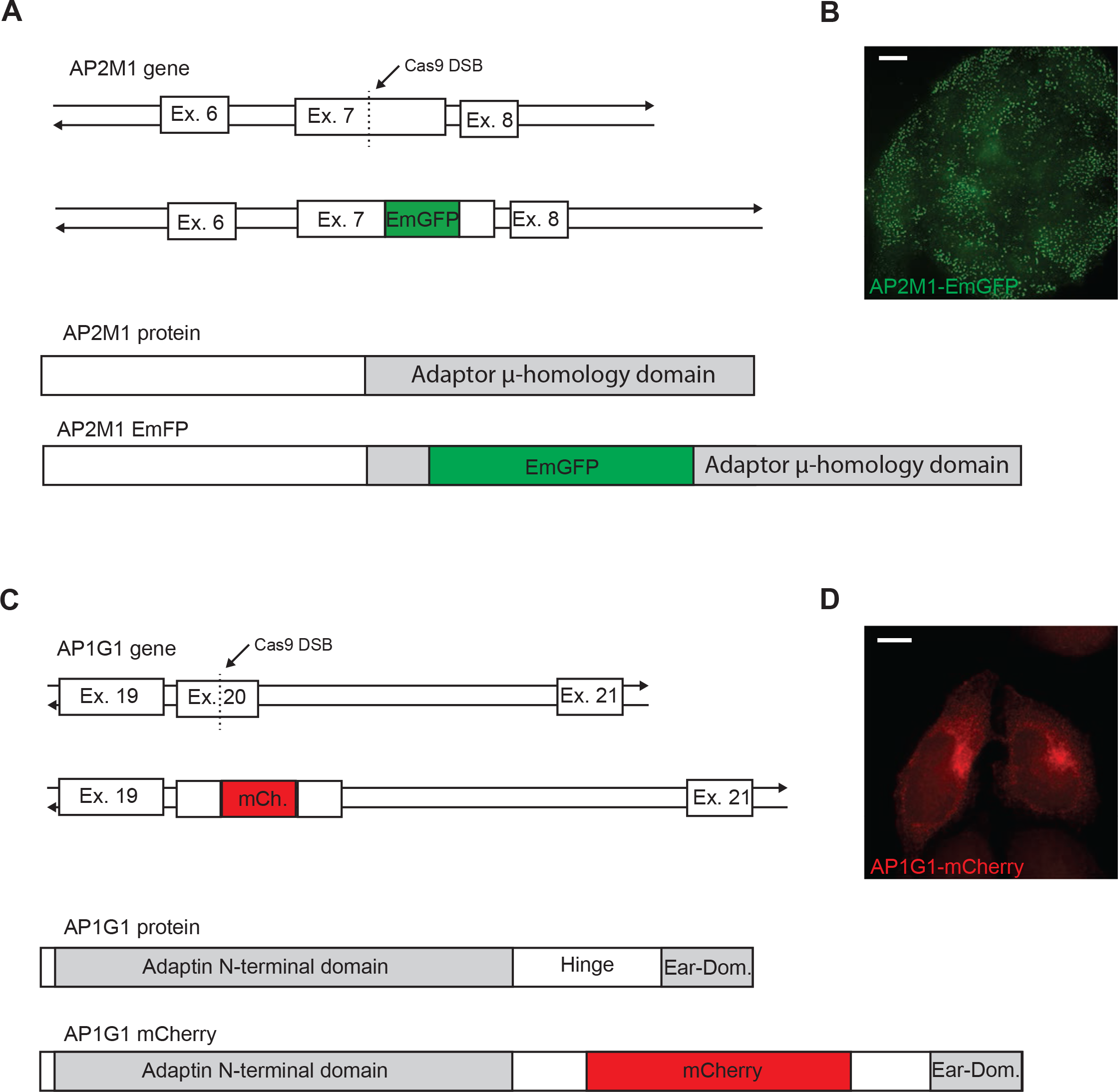
Internal tagging of AP2M1 and AP1G1. (A) To create an internal AP2M1-EmGFP fusion, a gRNA and PAM site was selected in exon 7 of the gene to generate an in-frame insertion of EmGFP into the C-terminal μ-homology domain of the protein when expressed. (B) Following isolation by flow cytometry, an EmGFP positive mixed population showed clear punctate plasma membrane EmGFP signal characteristic of endogenous AP2M1. (C) To tag AP1G1 with mCherry, a gRNA and PAM site was selected in exon 20 of the gene in order to place mCherry within the flexible hinge region of the expressed protein. (D) mCherry signal alone was at the limit of detection so the signal was amplified with an antibody against mCherry, revealing tubulovesicular perinuclear staining characteristic of endogenous AP1G1 Scale bars equal 10 μm.

## Discussion

NHEJ-mediated knock-in is known to function well in a wide variety of cell lines (Maresca et al., 2013, Lackner et al., 2015, Suzuki et al., 2016, Schmid-Burgke et al., 2016, Sawatsubashi et al., 2017). Indeed, we have achieved comparable results in HeLa, HEK293-T and NRK cells suggesting that our approach will be broadly applicable to mammalian model cell systems. In the current study we chose to tag loci with fluorescent protein tags as this represents by far the most widely used approach for determining protein localisation and carrying out affinity isolation. However, there is no inherent restriction in the exogenous DNA sequence encoded in the donor beyond recovery of edited cells. We suggest that this could be achieved through addition of antibiotic resistance cassettes when tagging at the C-terminus. Otherwise, this represents a problem common to all currently available genome editing approaches necessitating fluorescent tag use, an application which we have shown CHoP-In to greatly facilitate.

The current study suggests that N-terminal tagging is the optimal approach due to indel generation in other alleles when targeting the open reading frame. This limitation can likely be overcome however, either by working in a haploid or near haploid cell line such as HAP1 as previously demonstrated for NHEJ-based editing (Lackner et al., 2015), or alternatively by screening clonal cell populations for either multiple targeting events or the presence of unedited alleles. In fact, the decision to work in HeLa cells, with their polyploid nature, likely increased the likelihood of recovering alleles harbouring indels. Note also that when generating an internally tagged gene, although the probability of generating mutant alleles remains, it is likely that nonsense mediated decay would prevent the production of a truncated protein (Popp & Maquat, 2016).

As well as speeding up the generation of individual edited cell lines, CHoP-In has an inherent scalability, being based upon readily synthesised oligonucleotides. This feature of the approach will be particularly useful for following up uncharacterised proteins identified via genetic and proteomic screens (Kozik et al., 2013; Navarro Negredo et al., 2018). Validation and characterisation of these screening “hits” is often frustrated by overexpression artefacts and a lack of reliable antibodies for the analysis of endogenous proteins. Of particular concern for those interested in processes occurring at the subcellular level is the common observation of overexpression leading to aberrant localisation (e.g. Hirst et al., 2015). As such, the inherent speed and versatility of our approach will be useful to those wishing to accelerate the generation of multiple cell lines expressing tagged proteins at their endogenous levels and under endogenous control.

## Materials & Methods

### Cell culture

HeLa cells were cultured in RPMI 1640 (Sigma Aldrich) supplemented with 10% foetal calf serum, 2 mM L-glutamine, 100 U/ml penicillin and 100 μg/ml streptomycin.

### Antibodies

Anti-Rab5C (Ab199530, Abcam), Anti-red, for detection of mCherry, (5F8 Chromotek), Anti-ATP6V1G1 (16143-1-AP, Protein Tech Europe), Anti-EF2 (C14, Santa Cruz), Anti Lamp-1 (H4A3, Abcam), Alexafluor 568 anti-rat (Thermo Fisher). HRP-conjugated secondary antibodies (Sigma Aldrich).

### Genome editing by CHoP-In

*Streptococcus pyogenes* Cas9 and gRNAs were expressed from the pX330 plasmid (Addgene #42230) (Cong et al., 2013). Primers for generating CHoP-In integration fragments were designed as outlined in the current study. gRNAs were selected using the GPP web portal (Broad Institute), all gRNA and primer sequences are shown in supplemental table S1. Integration fragments were generated by PCR using KOD polymerase (Merck) from standard vectors pUC19-EmFP, pmCherry-N1. For each integration fragment the product of 5 × 100 μl PCR reactions was pooled and purified by ethanol precipitation prior to resuspension in 0.1 × TE at a concentration of approximately 2 μg/μl. HeLa cells grown on 9 cm plates were transfected using either lipofectamine 2000 (Thermo Fisher) or HeLa Monster (Mirus) according to manufacturers’ instructions. Cas9/gRNA expression plasmid and integration cassette were transfected at 1:1 ratio by mass. 48-72 hours post-transfection successfully edited cells were enriched by flow cytometry sorting and subsequently cultured either as a mixed population or further diluted to generate monoclonal cell lines.

### Genomic DNA isolation and sequencing

Genomic DNA was isolated from approximately 1 × 10^6^ cells using the High Pure PCR template preparation kit (Roche) according to manufacturers’ instructions. The region around the CRISPR/Cas9 lesion was amplified using primers outlined in supplemental table S1. PCR products were cloned into pCR-Blunt vector (ThermoFisher,), transformed into *E. coli* and resulting colonies screened by colony PCR for the presence of an integration event. Positive colonies were amplified, and recovered plasmids analysed by sanger sequencing using M13 reverse and T7 forward primers.

### Magic Red colocalisation

Magic Red cathepsin B substrate (Immunocytochemistry Technologies) was prepared as previously described (Bright et al., 2016). Cells were grown on PeCon glass coverslips or MatTek glass bottomed dishes for live cell imaging. Cells were incubated at 37°C in a 5% CO_2_ incubator for at least 2 minutes in the presence of the cathepsin substrate before being transferred to the microscope for imaging.

### Transferrin colocalisation

Cells, seeded onto glass cover slips, were incubated for 15 minutes at 37°C in the presence of Alexa-fluor 555 labelled human transferrin (Thermo Fisher) (25 μg/ml) in serum free media. Cells were then washed in ice cold PBS prior to fixation for 10 minutes in 4 % (w/v) paraformaldehyde in PBS and processing for fluorescence microscopy.

### Immunofluorescence staining

Cells, seeded onto glass coverslips, were washed in ice cold PBS and fixed for 10 minutes in 4% w/v) in PBS. Following fixation, cells were permeabilised in 0.1% Triton X-100 in PBS for 15 minutes at room temperature before blocking for 1h in 20% foetal calf serum in PBS. Primary and secondary antibody incubations were carried out sequentially at room temperature for at least one hour in 20% foetal calf serum in PBS.

### Fluorescence microscopy

For fixed-cell samples, coverslips were mounted onto glass slides in ProLong Diamond antifade reagent with DAPI (Thermo Fisher). Cells were imaged on a Zeiss Axio Imager upright microscope under a 63 × 1.4 NA Plan Apochromat oil immersion objective. Live cell imaging was carried out on a Zeiss confocal microscope equipped with an incubated stage.

### Immunoblotting

Immunoblotting was conducted as previously described (Navarro Negredo et al., 2018). Briefly, cells were lysed in SDS buffer (2.5% SDS, 50 mM Tris, pH 8.0) before boiling in NuPAGE LDS sample buffer supplemented with 0.1M DTT. SDS-PAGE was performed on NuPAGE 4–12% Bis–Tris gels with NuPAGE MOPS SDS Running Buffer (Life Technologies). Following transfer of proteins to nitrocellulose membranes, membranes were blocked with 5% non-fat milk in PBS with 0.1% (v/v) Tween-20 (PBS-T). Primary antibody incubations were carried out for at least one hour with appropriate horseradish peroxidase-conjugated secondary antibodies applied subsequently. Chemiluminescent detection of bound antibody was carried out using AmershamECL Prime Western Blotting Detection Reagent (GEHealthcare) and X-ray film (Kodak).

### Flow cytometry

2-5 × 10^6^ cell were trypsinised, washed with PBS and resuspended at a concentration of 10^6^/ml in PBS. Sorting was carried out on a BD FACSMelody cell sorter. Positive cells, gated at a fluorescent intensity above all events seen in a control population, were collected and seeded into 25 mm plates for expansion to a mixed population.

---

**Supplemental figure S1.**
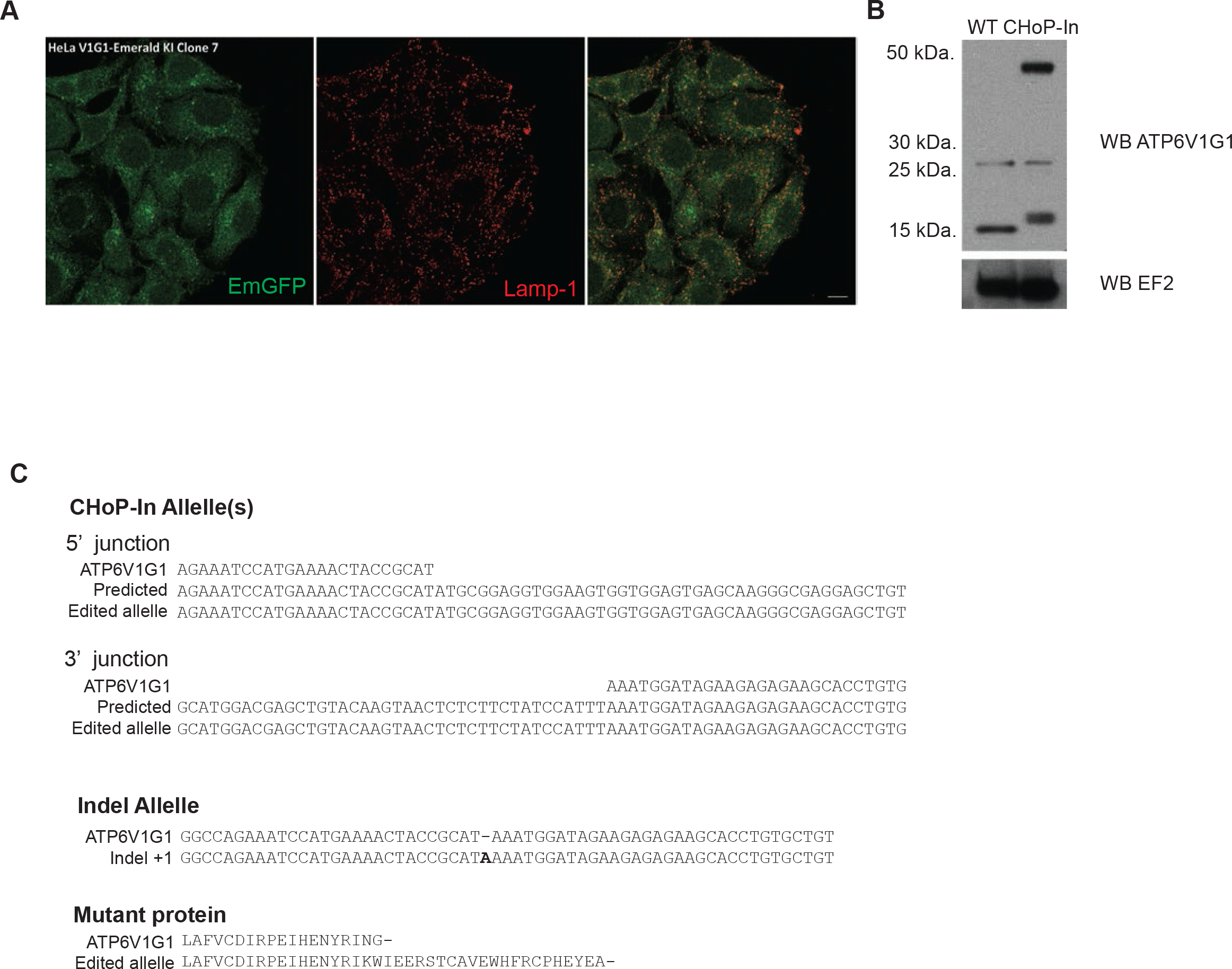
Analysis of a monoclonal ATP6V1G1-EmGFP cell line. (A) A clonal cell line (clone 7) isolated from the ATP6V1G1-EmGFP mixed population showed correct localisation of EmGFP signal (green) assessed by colocalisation with the endolysosomal marker LAMP-1 (red) (scle bar equals 5 μm). (B) Immunoblot with an antibody against ATP6V1G1 revealed that in addition to the higher molecular weight band corresponding to the ATP6V1G1-EmGFP fusion, the band corresponding to the unedited wild type protein was replaced by one of slightly higher molecular weight. (C) Sanger sequencing confirmed at least one correctly edited allele as well as a further allele harbouring a one bp indel, generating a frame-shift mutation and extending the reading frame of ATP6V1G1 at this allele accounting for the shift in molecular weight observed by immunoblotting.

**Supplemental figure S2.**
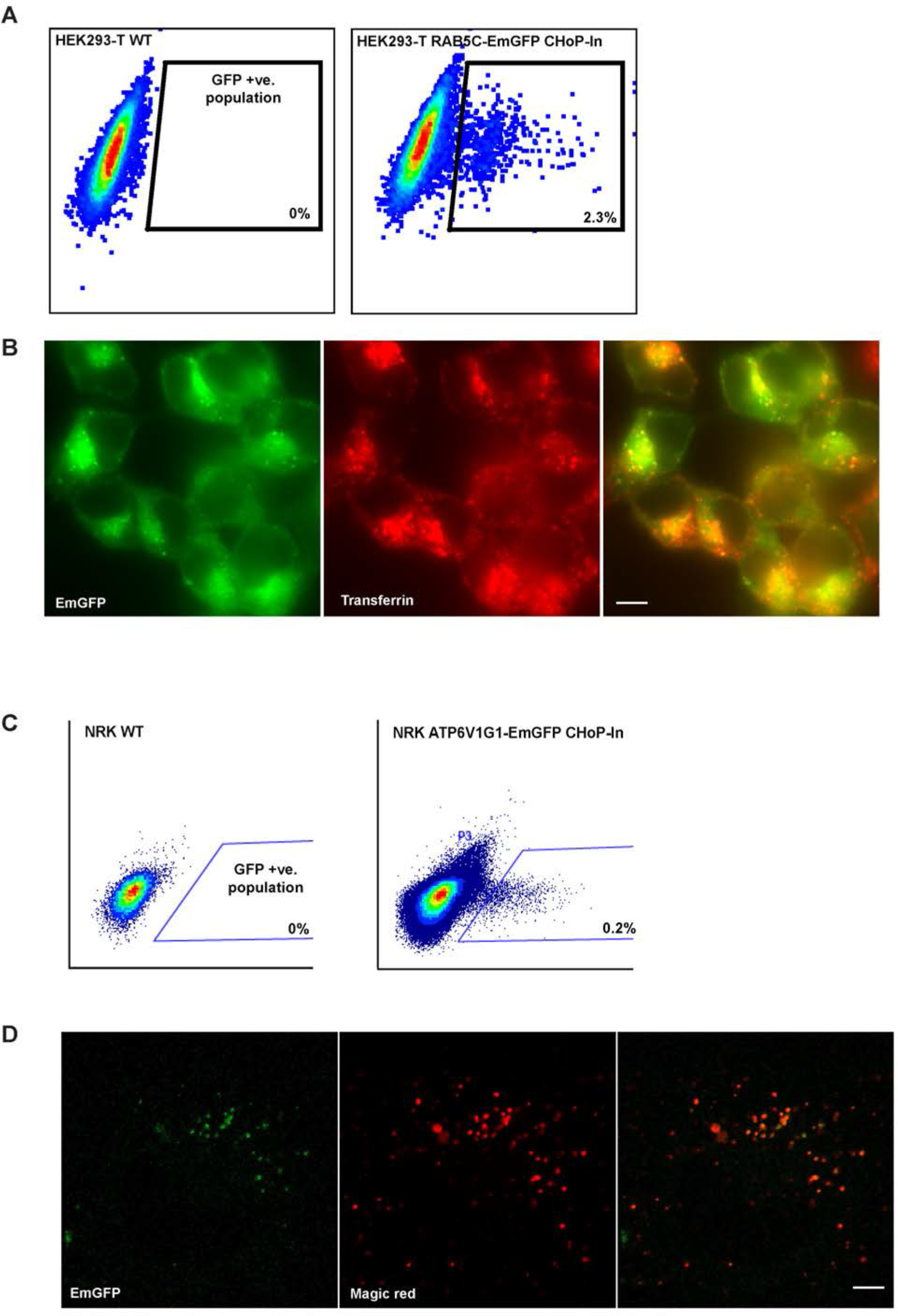
CHoP-In editing in other cell types. HEK293-T cells were edited using CHoP-In to express an EmGFP RAB5C fusion from its endogenous locus and NRK cells were edited to express an ATP6V1G1-EmGFP fusion. (A) EmGFP-Rab5C expression was assessed in transfected HEK293-T cells by flow cytometry. WT cells are untransfected HEK293-T. (B) EmGFP positive HEK293-T were assayed for correct localisation of EmGFP-Rab5C fusion by colocalisation with endocytosed transferrin. (C) CHoP-In edited NRK cells were assessed and sorted by flow cytometry. WT cells are untransfected NRK. (D) Correct localisation of ATP6V1G1-EmGFP was assessed by colocalisation of EmGFP signal with the endo-lysosomal marker magic red.

**Supplemental figure S3.**
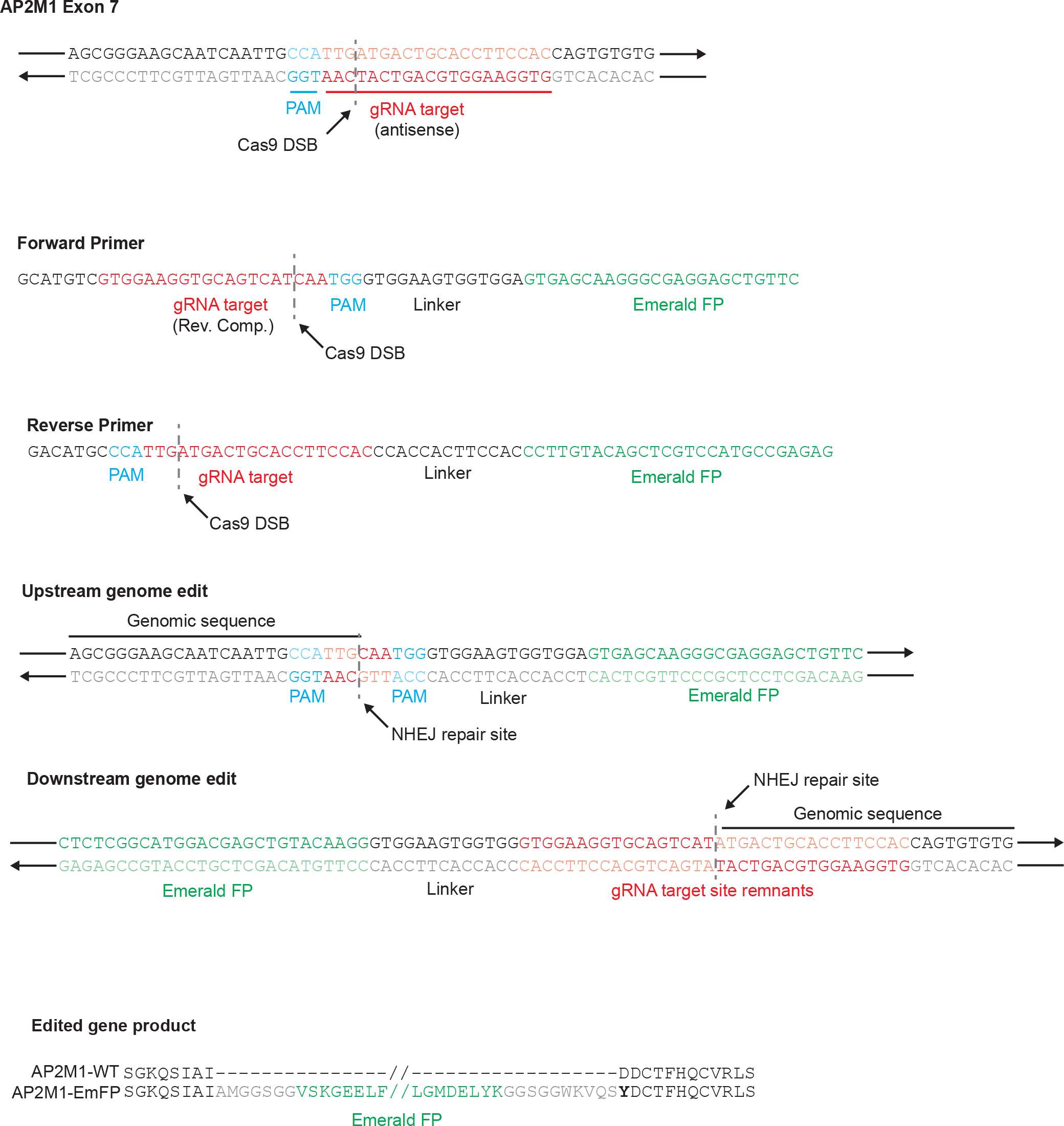
ChoP-In strategy for creating internal EmGFP fusion of AP2M1. Detailed description of the CHoP-In editing strategy used to create the internal AP2M1-EmGFP fusion.

**Supplemental figure S4.**
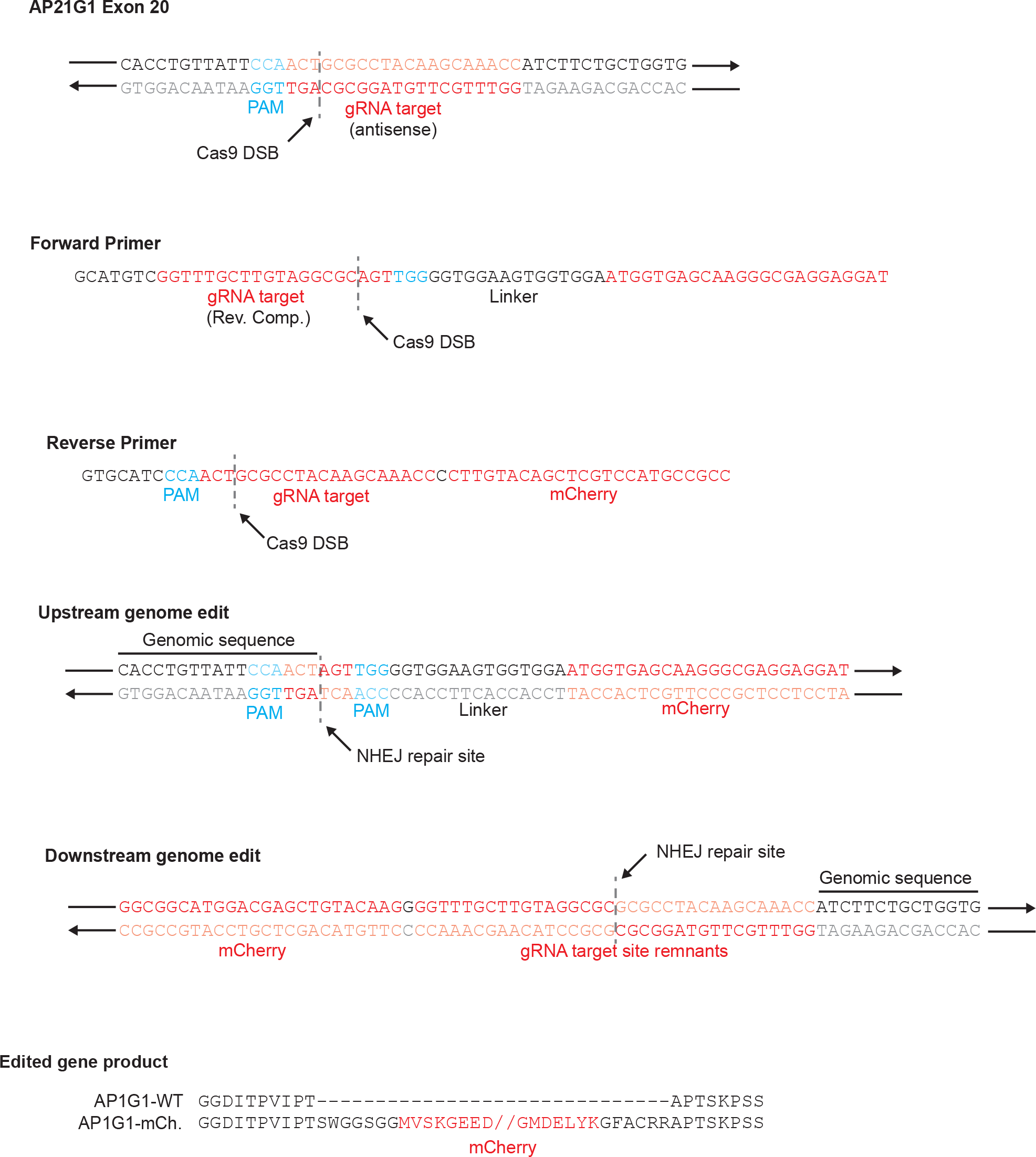
CHoP-In strategy for creating internal mCherry fusion of AP1G1. Detailed description of the CHoP-In editing strategy used to create the internal AP1G1-mCherry fusion.

**Supplemental Table S1.**
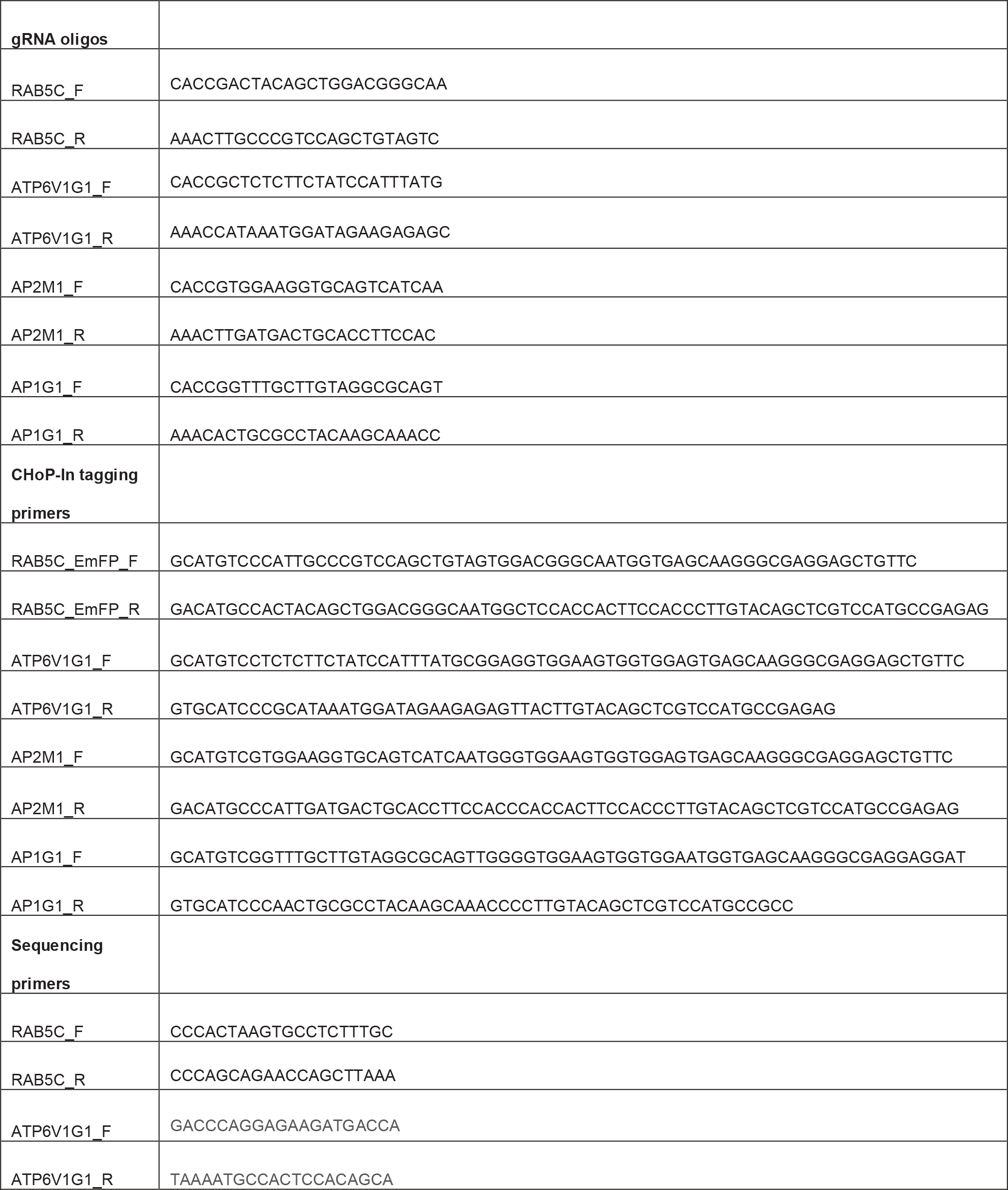
Oligonucleotides used in the current study.

